# Immunodominant influenza epitope GILGFVFTL engage common and divergent TCRs when presented as a 9-mer or a 15-mer peptide

**DOI:** 10.1101/2022.07.11.499638

**Authors:** Keshav Bhojak, Vasumathi Kode, Coral M. Miriam, Kayla Lee, Athulya Ramesh, HV Sudheendra, Papia Chakraborty, Amitabha Chaudhuri

## Abstract

Antigen-specific T-cells are a powerful modality for treating cancer and other life-threatening viral and bacterial diseases. Technologies to identify and expand antigen-specific T-cells rapidly can shorten the time, and lower the cost of treatment. In this regard, screening short overlapping peptides to identify antigen-specific T-cells in *ex vivo* T-cell activation assay is becoming routine. Screening assays typically use 15-mer peptides to stimulate patient-derived, or healthy peripheral blood mononuclear cells to activate T-cells and identify expanded TCRs by next generation sequencing. Previous studies comparing the kinetics of T-cell activation using a 9 and a 15-mer peptide versions of a CMV immunodominant epitope demonstrated that 15-mer peptides induced CD8 T-cell activation at a slower kinetics reaching a lower magnitude compared to 9-mer peptides. The fact that 9-mer peptides are an optimal fit for the MHC class-I binding groove could explain this difference, with the 15-mer peptide requiring additional proteolytic processing before binding to the deeper binding groove of class-I MHC. Alternatively, the delay in kinetics and magnitude can result from the activation of a wider diversity of TCRs engaging novel epitopes generated from the 15-mer peptide whose activation profile may be different from the profile of TCRs that normally respond to the 9-mer immunodominant epitope. We sought to address these two possibilities by comparing T-cell engagement to the HLA-2-restricted GILGFVFTL epitope presented as a 9-mer, or a 15-mer peptide and analyzing CDR3 expansion as a measure of T-cell engagement diversity. This approach also addressed an important question as to whether optimal TCRs could be missed using a 15-mer peptide used routinely in screening assays.

## Introduction

Discovery of immunodominant epitopes from infectious agents, cancer and autoimmune diseases has revealed immunological underpinnings of protective/destructive immunity, paving the way for novel interventional therapies (1, 2). Therapeutic vaccines comprising of immunodominant epitopes induce strong CD8 and/or CD4 T-cells to generate long-lasting immunity against infectious diseases, including cancer (3-6). However, such protective immunity is modulated by many different factors such as antigen dose, strength of TCR signaling, duration of activation and quality of T-cells generated in response to the antigen (7-9). Studies using immunodominant epitopes from influenza, HIV and other infectious agents have demonstrated that of the many immunodominant epitopes identified by screening affected donors, only a handful induced cytolytic CD8 T-cells that correlated with protection (10-12). Therefore, there is a need to screen more antigens to identify epitopes that can be developed into therapeutic vaccines.

Efforts to discover immunogenic epitopes from different sources are performed routinely by high throughput T-cell activation assays (13-15). In these assays, peptides are added from outside to donor PBMCs and their immunogenicity is determined over a period of time by quantitating markers of T-cell activation. For an exploratory screen, 15-mer overlapping peptide pools covering the length of the antigen are synthesized and mixed with donor PBMCs usually at 10μM and activation monitored by IFN-γ expression or expression of other activation markers by T-cells (16).

For CD8 T-cell activation, 9-11 mer peptide length is optimal for binding MHC class-I binding groove. However, 15-mer peptides also activate CD8 T-cells as a result of peptide trimming to achieve a close fit within the MHC class-I binding groove (7, 17). A comparison between T-cell activation by 9-mer and 15-mer peptides revealed slower kinetics of activation by 15-mer peptides and requirement of a higher dosage of the antigen for activation (17). Antigen dosage determines the quality of T-cells. Both preclinical models and human studies show that in TB and *Leishmania major*, for example, a higher antigen dose causes activation-induced cell death (AICD) of T-cells, selecting cells that may be functionally inferior in clearing infection and generating a robust protective immunity (17-21). By contrast, new-born and elderly population responded less well to low antigen dosage vaccination than individuals outside these age groups indicating lack of optimal T-cell repertoire diversity that may not have formed yet in the new-born and is lost in the aged population (9).

Our goal in this study was to determine correlation between TCR expansion and the pattern of CD8 T-cell activation by 9 and 15-mer peptides when used at a low and high antigen dosage. We examined expansion of TCRS by using the 9-mer immunodominant flu epitope GILGFVFTL and created a 15-mer version LTKGILGFVFTLTVP by adding three amino acids to either end of the 9-mer peptide. We used both the peptides at a low and a high dosage and analyzed the kinetics and magnitude of CD8 T-cell activation along with the expansion of CDR3s at different timepoints. The results confirmed the earlier findings that the 15-mer peptide induced CD8 T-cell activation at a slower kinetics, but reached a similar magnitude as the 9-mer peptide at a later timepoint. However, the longer peptide only activated CD8 T-cells when used at 10μM, but not at 100nM concentration. This is in contrast to the 9-mer peptide that activated T-cells at both concentrations. The pattern of expansion of CDR3s however, demonstrated sensitivity of CD8 T-cells to the 9-mer peptide when used at a high and low antigen dosage. Some of the CDR3s expanded by the 15-mer peptide in this study are bonafide binders of the 9-mer epitope suggesting that overlapping 15-mer peptides used in high throughput screening are likely to identify CDR3s that are cognate binders of 9-mer immunodominant epitopes.

## Results

### CD8 T-cell activation by the GILGFVFTL peptide

We analyzed the kinetics and magnitude of intracellular IFN-γ expression by CD8 T-cells in response to the 9-mer GILG peptide at two different concentrations in four different HLA-A.02 donors. As shown, 10μM peptide induced robust IFN-γ activation (10-30% IFNγ^+^ CD8 T-cells) in all donors except D12299 on day-7, decreasing in magnitude over time (Figure 1A-D). There was variability in the kinetics of IFN-γ activation between donors (Figure 1A-D). For example, in donor D17843, maximal activation was observed on day-7 followed by a sharp decline in the signal by day-14 (80% decline) (Figure 1B), whereas in D089 and D384 activation was sustained until day-14 and decreased by day-21 (Figure 1C-D). In D12299 by contrast, 10μM peptide induced a weaker response compared to 100nM peptide (Figure 1A) contrasting sharply with other donors where both peptide concentrations generated robust T-cell activation (Figure 1B-D).

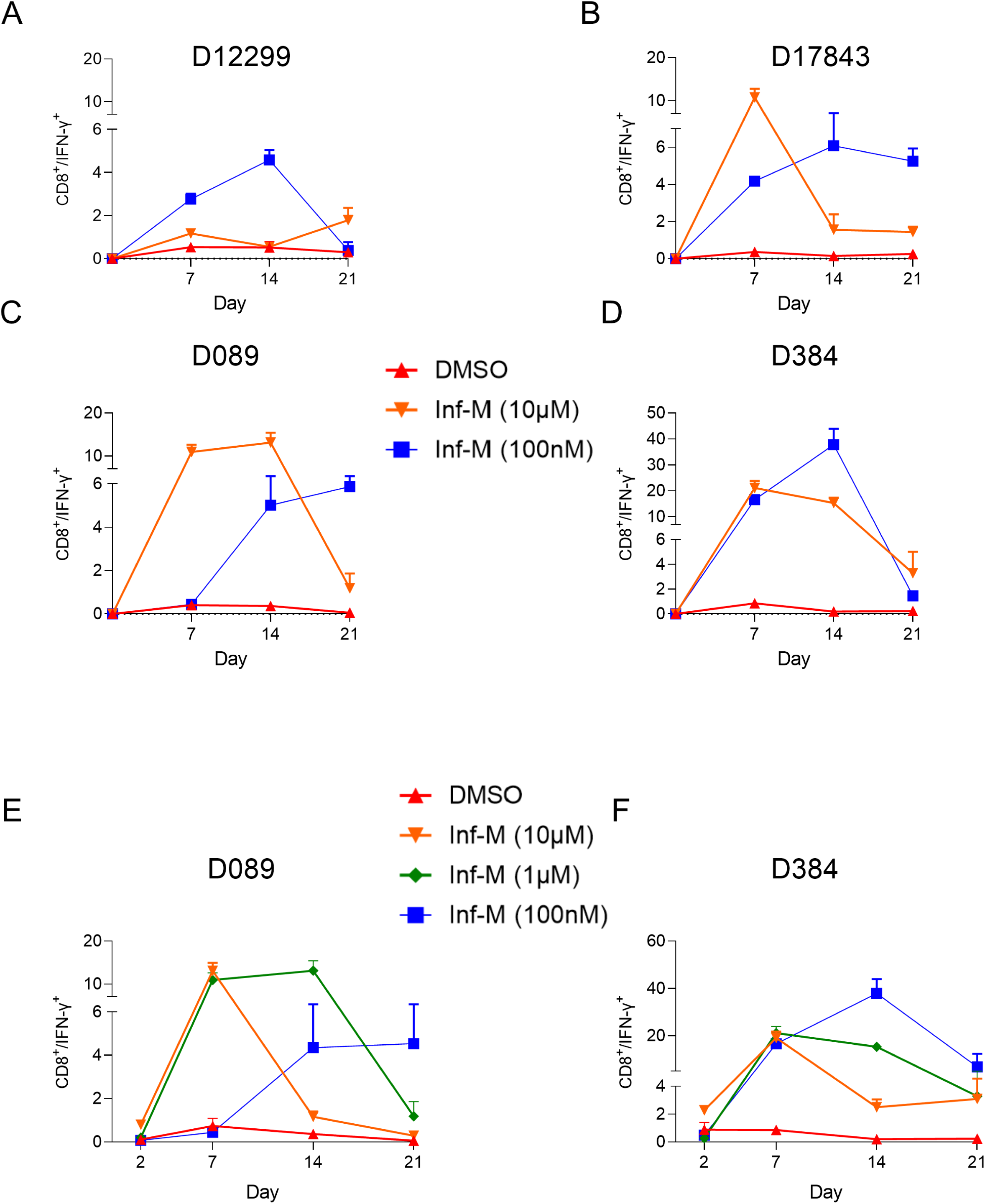

Donor to donor variability in response to immunodominant epitopes have been observed in other studies and is often associated with susceptibility, or resistance to infectious diseases (22, 23). We selected donors D089 and D384 to further investigate differences in T-cell activation kinetics at an earlier timepoint in the presence of different peptide concentrations (Figure 1E-F). In D089, 10μM peptide induced weak IFN-γ activation at day-2 (<1%) and undetectable activation by 1μM and 100nM peptide. Maximal activation was reached at day-7 following which activation declined sharply in 10μM peptide, whereas activation was sustained till day-14 by 1μM peptide (Figure 1E, orange and green lines). In contrast, 100nM peptide induced T-cell activation at a slower kinetics (Figure 1E blue line) reaching maximal activation on day-14, which was 50% compared to what was reached at higher peptide concentrations. Compared to donor D089, however, the pattern of T-cell activation in donor D384 was different. Robust activation >2% was detected at day-2 in the presence of 10μM peptide, indicating strong pre-existing immunity against influenza in this donor. The decline in activation was rapid at higher peptide concentrations and was sustained for a longer time period at 100nM (Figure 1F). Taken together, in both donors, at higher peptide concentrations there was a rapid burst in CD8 T-cell activation followed by a kinetics of decline that was inversely proportional to the peptide concentrations. In addition, compared to donor D089, donor D384 was more responsive to the GILG peptide, exhibiting 8-fold higher T-cell activation at 100nM peptide (5% vs 40%) respectively.

Given donor D384’s strong response to the GILG peptide, we modeled the magnitude and kinetics of CDR3 expansion to the 9-mer and a 15-mer version of the GILG peptide (core 9-mer epitope, flanked on either side by three amino acids) in this donor.

As expected, the 9-mer peptide induced rapid CD8 T-cell activation compared to the 15-mer peptide (Figure 2A). At 10μM, the 9-mer peptide induced maximal CD8 T-cell activation by day-2 (11%), which declined to the baseline by day-21 (Figure 2A, green squares). Compared to 10μM peptide, 100nM peptide induced five-fold lower CD8 T-cell activation at day-2 (Figure 2A). However, by day-7 and day-14, 100nM peptide surpassed the activation magnitude of 10μM peptide by ∼3-fold (30% vs 11%) (Figure 2A, blue and green lines). The 15-mer peptide showed robust activation similar to 100nM 9-mer peptide only at 10μM, with minimal activation at 100nM peptide concentration (Figure 2A, red and magenta lines). The results confirmed previous observation (17) that the kinetics of CD8 T-cell activation was faster in the presence of the 9-mer peptide than the 15-mer peptide and comparable magnitude of activation by the 15-mer peptide occurred at 60-fold higher concentration compared to the 9-mer peptide. Another interesting difference between the shorter and the longer peptides was that the 9-mer peptide failed to preserve the activation state of T-cells over an extended time period compared to the 15-mer peptide. For example, by day-21, 10μM or 100nM 9-mer peptide retained ∼10% of its maximal activation, whereas 10μM 0f the 15-mer peptide retained 50% of its maximal activation (30% to 15%) although both peptides reached peak activation on day-14. The decline in CD8 T-cell activation after day-7 in the presence of 10μM GILG peptide (Supplementary Figure 1A) appears to be due to a decrease in the absolute number of CD8^+^ T-cells (Supplementary Figure 1B) and increase in the expression of granzyme-B (GZMB) (Supplementary Figure 1C) a hallmark of activation-induced cell death (24, 25).

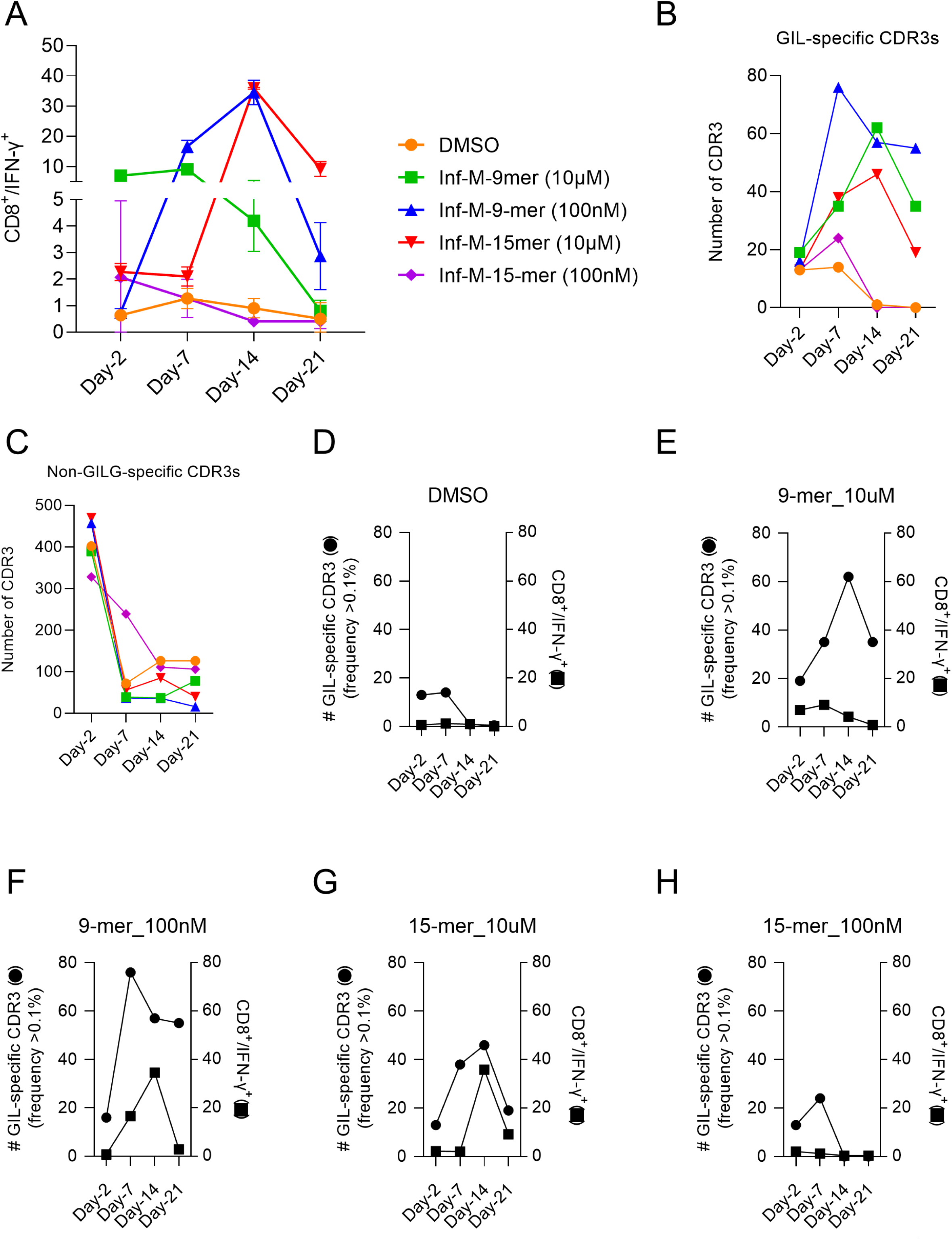

Although processing of the 15-mer peptide before binding class-I HLA can explain the delay in T-cell activation kinetics, the preservation of the T-cell activation state in the presence of the longer peptide (50% residual activity for the 15-mer vs 10% for the 9-mer peptide (Figure 2A) raised the possibility that besides CD8 T-cell death, changes in the repertoire structure and the clonal expansion of CDR3s may be additional contributing factors.

### Changes in CD8 T-cell repertoire in response to 9-mer and 15-mer peptides

Immune epitope database (IEDB) and VDJDB databases list unique TCR-β sequences against the 9-mer GILGFVFTL peptide (GIL-peptide). There are no TCR-β for the 15-mer peptide LTKGILGFVFTLTVP in these databases. We analyzed changes in GIL-specific CDR3-β repertoire separately from CDR3s that are not reported in public databases to be GILG-specific (non-GILG-specific CDR3s) by the two peptides at different timepoints.

We detected 3830 unique CDR3s over 0.1% frequency in this study (Supplementary Table 1), of which 143 were GILG-specific (Supplementary Table 2). Increase in GILG-specific CDR3 numbers correlated with T-cell activation kinetics (Figure 2B). Lack of T-cell activation correlated with the expansion of a smaller number of unique CDR3s (14 - 24 unique CDR3s in DMSO (orange circle) and 100nM 15-mer peptide (purple diamond) respectively) (Figure 2B). In contrast, strong T-cell activation correlated with 50-70 GILG-specific CDR3s in different treatments (Figure 2B). The number of GILG-specific CDR3s detected in the presence of the 9-mer peptide was higher (65 CDR3s) compared to the 15-mer peptide (46 CDR3s) (Figure 2B) even though magnitude of CD8 T-cell activation was comparable between 9 and 15-mer peptides (Figure 2A). The expansion of non-GILG-specific CDR3s however, showed an opposite trend in which CDR3 numbers declined under all treatment conditions with the highest number of CDR3s detected at day-2 (Figure 2C).

Next, we analyzed changes in the frequency of GILG-specific CDR3 with the kinetics of CD8 T-cell activation to examine relationships between the two events. We observed that 10μM of the 9-mer peptide induced peak activation (day-7) before reaching the peak of CD8 T-cell expansion that occurred later (day-14) (Figure 2E), whereas the same antigen at 100nM induced peak CDR3 expansion earlier (day-7), before reaching peak CD8 T-cell activation (day-14) (2F). By contrast both CD8 T-cell activation and CDR3 expansion occurred at the same time in the presence of 10μM 15-mer peptide (Figure 2G).

Next, we examined top-10 maximally expanded CDR3s under different treatment conditions to identify common and divergent CDR3s across different treatments. GILG-specific CDR3s showed robust expansion under conditions that induced strong CD8 T-cell activation, but failed to expand under conditions that did not induce activation (Figure 3A). Two GILG-specific CDR3s CASSIRSSYEQYF (blue circle) and CASAHPRASGSTDTQYF (orange circle) reached ≥4% frequency in the presence of 10μM or 100nM of the 9-mer peptide and 10μM 15-mer peptide suggesting that these CDR3s recognize the immunodominant GILGFVFTL epitope and that the 15-mer peptide is processed to generate the immunodominant epitope. Interestingly, CASSYNTEAFF (yellow circle) was strongly expanded in the presence of 10μM 15-mer peptide, reaching a frequency of 16%, although its level of expansion was ≤4% in the presence of the 9-mer peptide suggesting that this CDR3 may be a private CDR3 or a CDR3 that recognizes an epitope overlapping with the 9-mer GILG epitope. A second CDR3, CASSARSSTEQYF (gray circle) was 3% expanded by 10μM 9-mer or 15-mer peptide and 9% expanded at 100-fold lower concentration of the 9-mer peptide suggesting that it is a dosage-sensitive CDR3. As expected, higher overlap in GILG-specific CDR3s were detected between treatments that led to strong CD8 T-cell activation (Figure S2A).

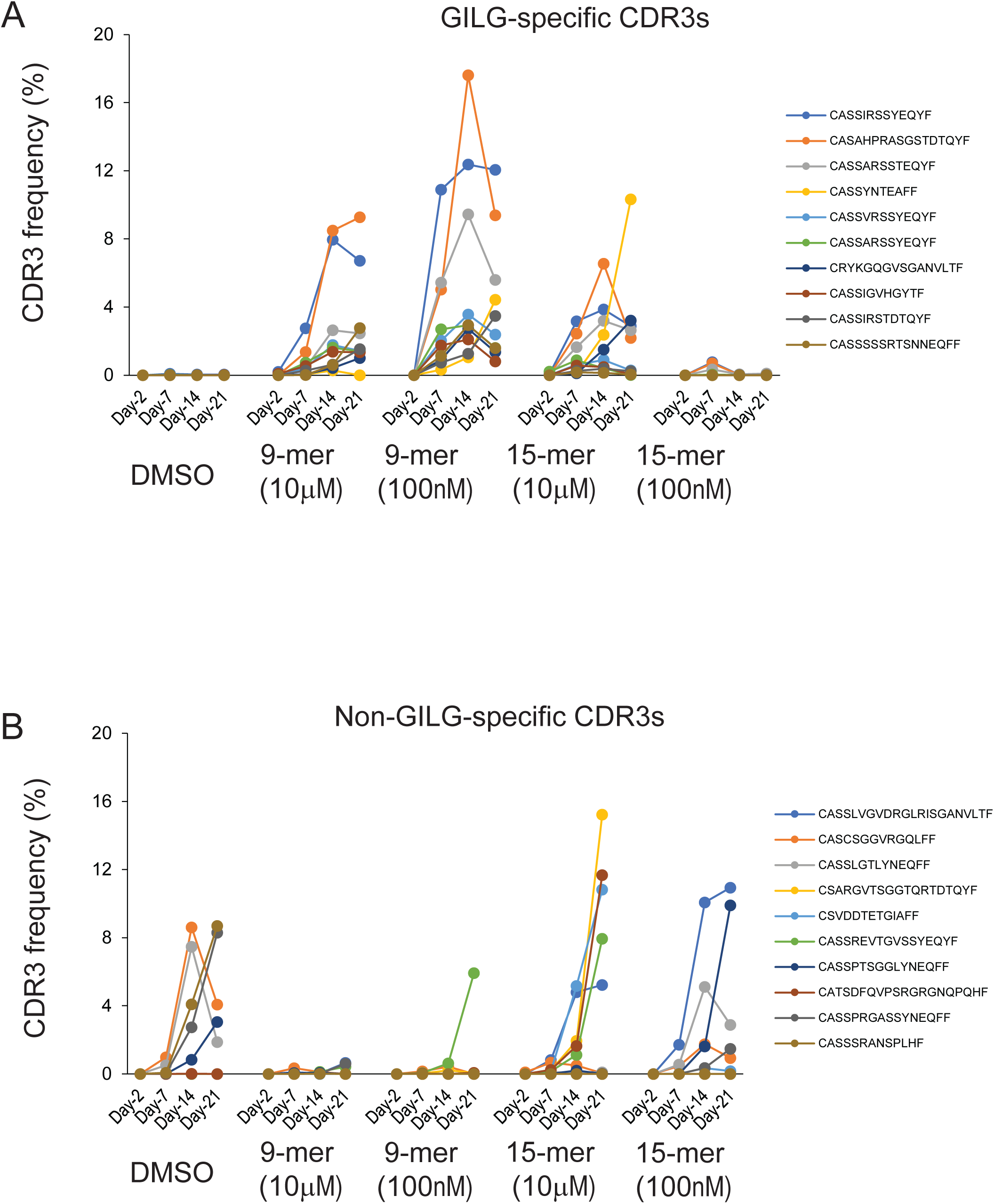

The expansion of non-GILG-specific CDR3s under different conditions was particularly interesting. Contrary to the expansion of GILG-specific CDR3s, the frequency of non-specific CDR3s remained stable under conditions that failed to induce CD8 T-cell activation (DMSO or 100nM 15-mer peptide). As expected, under conditions of CD8 T-cell activation by the 9-mer peptide, few CDR3s were expanded (Figure 3B). Surprisingly, non-GILG-specific CDR3s were expanded by 10μM 15-mer peptide, even though there was robust CD8 T-cell activation (Figure 3B), suggesting that the 15-mer peptide may be engaging CDR3s from CD4 T-cells or recognizing novel epitopes generated from the processing of the 15-mer peptide. Five of the ten non-GILG-specific CDR3s expanded in DMSO were also expanded to various levels in the presence of 100nM of the 15-mer peptide suggesting non-specific expansion. Few non-GILG-specific CDR3s were common between treatments that led to strong CD8 T-cell activation (Figure S2B). The CDR3β CASSREVTGVSSYEQYF (green circle) showed >5% expansion in the presence of 100nM 9-mer and 10μM 15-mer peptide, but not in the presence of 10μM 9-mer peptide suggesting that it may be a novel CDR3 recognized by the GILGFVFTL peptide at a lower dosage and not reported in the IEDB database. Finally, two CDR3βs of, CSARGVTSGGTQRTDTQYF (yellow circle) and CATSDFQVPSRGRGNQPQHF (brown circle) were expanded to over 10% only in the presence of 10μM of the 15-mer peptide suggesting that these CDR3s may be recognizing epitopes other than the GILG immunodominant epitope. Spectratype analysis at day-14 indicated that 15-mer peptides expanded a CDR3β of 24 amino acids, absent from 9-mer treatments (Figure S3). Taken together, these observations suggest that the longer 15-mer peptide expanded multiple GILG-specific CDR3s that were also expanded by the 9-mer peptide and a few CDR3βs specific to the longer peptide.

### V-J gene usage by 9-mer and 15-mer peptides

Next, we compared the V and J gene usage for the TCR-β chain in the presence of 9-mer and 15-mer peptides and analyzed their enrichment over time. The V genes 20-1 and 19 were detected at the baseline and both these genes were detected in the presence of different treatments (Figure 4A-F). TRBV19 present at baseline (Figure 4A) showed non-specific expansion in the absence (Figure 4B and 4F) or presence (Figure 4C-E) of T-cell activation, although the expansion reached above 60% under conditions of T-cell activation (Figure 4C-E). TRBV19 usage is common in adults carrying pre-existing TCRs against influenza-A virus (26). TRBV7-2, however, was expanded only under conditions of T-cell activation (Figure 4C-E) suggesting usage in functional TCRs, whereas TRBV13 was expanded only in DMSO and 100nM 15-mer peptide (Figure 4B and 4F). Expansion of TRBV6-6 (grey circle) occurred only in the presence of 10μM 15-mer peptide (Figure 4E) suggesting that this gene could be responsible for the generation of both GILG-specific and non-GILG-specific epitopes by the 15-mer peptide (Figure 4E). A differential expansion pattern for the J-gene was not observed, except TRBJ1-2, which was expanded only in the presence of 10μM 15-mer peptide (Figure S4).

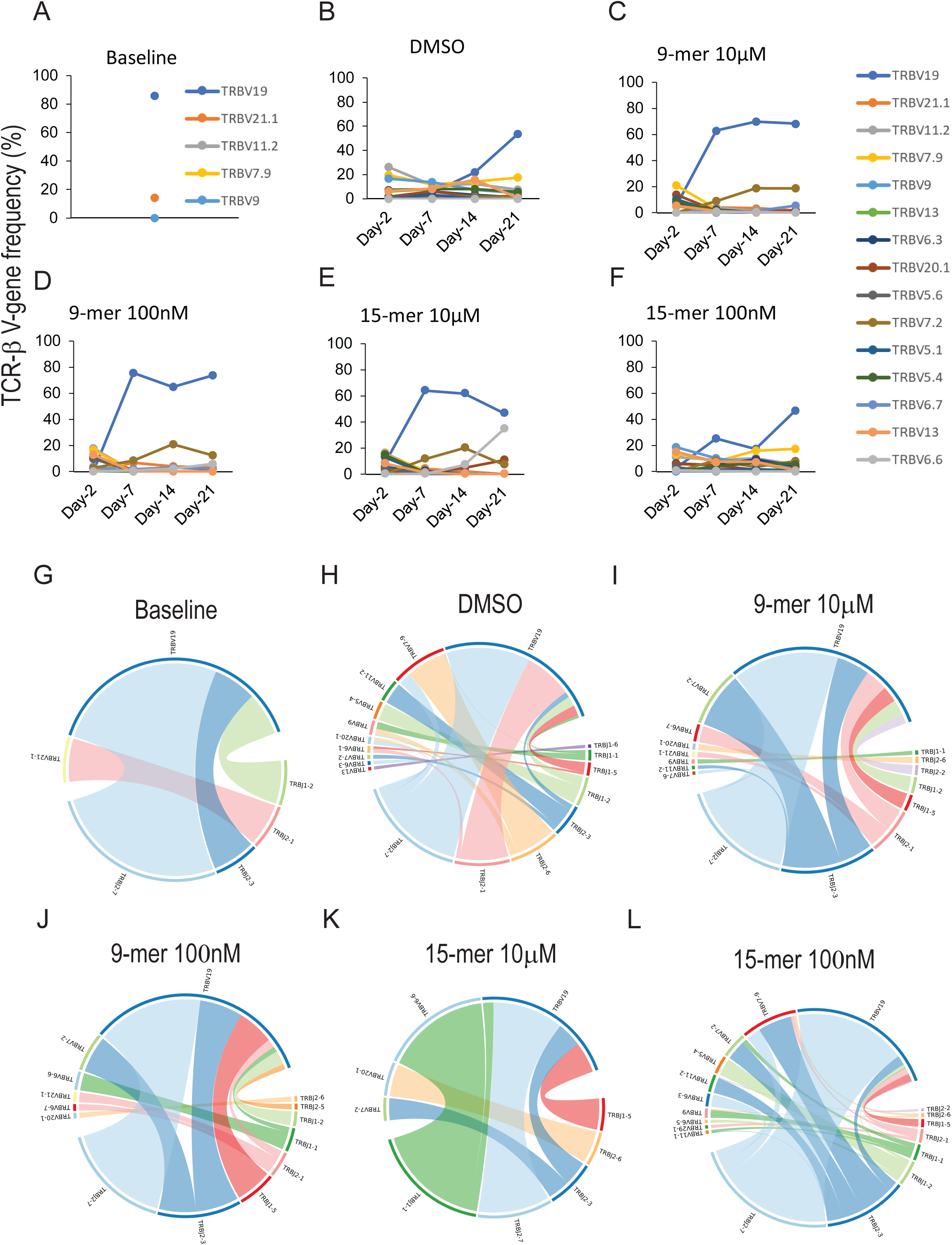

We observed enrichment of multiple V-J recombination events associated with response, or a lack of response to the peptides. For example, TRBV19 – TRBJ2-7 was dominant at baseline and was detected both under conditions of T-cell activation, or lack of activation (Figure 4G-L). However, TRBV19 - TRBJ2-3 recombination was present at baseline, minimally expanded in DMSO or 100nM 15-mer peptide (Figure 4G-H and L)), but expanded robustly under conditions of activation (Figure 4I-K). A second recombination event TRBV7-2 – TRBJ2-3 was not observed at baseline (Figure 4G), present at a low frequency in DMSO (Figure 4H), but expanded significantly in the presence of 9-mer and 15-mer peptides (Figure 4I-L) suggesting that this combination is associated with antigen presence. A highly expanded TRBV6-6 – TRBJ1-1 recombination event was observed in the presence of 10μM 15-mer peptide as expected from the specific expansion of TRBV6-6 noted earlier (Figure 4K). However, TRBJ1-2 specifically expanded by 15-mer 10μM peptide was not represented in the most frequent V-J recombination events associated with GILG-specific CDR3s (Figure 4K). Among the non-GILG CDR3s 10μM 15-mer peptide expanded TRBV20-1 – TRBJ2-3 specifically (Figure S5E).

### T-cell activation and CDR3 expansion by the CEF pool

Having determined the kinetics and magnitude of expansion of CDR3s in response to the 9-mer GILG peptide, we sought to examine the CDR3 expansion pattern when presented with a mixture of immunodominant epitopes, such as the CEF pool containing five HLA-A2 restricted 9-mer peptides, including GILGFVFTL. Individual peptides in the pool are present at 15nmole/peptide. Donor D384 was treated in triplicates with CEF pool in which each peptide is present at 100nM final concentration and a time course was performed in triplicate. CD8 T-cell activation profile was comparable in the three replicates (Figure 5A). The kinetics and magnitude of activation mimicked the response of D384 to 100nM of GILG peptide (Compare Figure 5A and 1F). Having an explicit knowledge of CDR3s expanded by pure 9-mer GILG peptide, we analyzed the expansion of CDR3s by the CEF pool and used the CDR3s as a marker to ascertain the contribution of the GILG peptide on the CDR3 repertoire of D384. The expansion of the top-6 GILG-specific CDR3s is shown in Figure 5B-D). These CDR3β chains were also expanded in the presence of 100nM pure GILG peptide. Five of the six CDR3βs were expanded over 1% in the presence of the CEF pool (Figure 5D) and were present at a low frequency at baseline or in DMSO, indicating antigen-specific expansion (Figure 5B-C). The peptide CASSSGLVSNTGELFF was detected at a frequency of 0.7% at baseline (Figure 5B), failed to expand in the presence of DMSO (Figure 5C) and expanded up to 14% in the presence of CEF pool. Interestingly, CASSIRSSYEQYF was expanded by 100nM GILG-peptide alone, (Figure 3B) and the CEF pool (Figure 5D). In both instances, maximal expansion was observed at day-14 although magnitude of expansion was higher in the presence of peptide alone (10%) than by the CEF pool (3%) (Compare 3B with 5D) suggesting competition between peptides to engage TCRs, when present in a pool. Other CDR3s detected in response to the individual peptide was not expanded in CEF pool, and vice versa, suggesting differences in the availability of a specific peptide when present in a pool vs in a pure form. Next, we analyzed V-J recombination events in the presence of the CEF pool to determine if the recombination frequencies recapitulated what was observed with the pure GILG peptide (Figure 5E-F). In DMSO, TRBV9 – TRBJ2-2 was the most common recombination event (Figure 5E), whereas CEF treatment expanded TRBV19 – TRBJ2-7 in two of the three replicates and TRBV19 – TRBJ1-5 and TRBV7-2 – TRBJ2-3 in the third (Figure 5F). In the presence of GILG peptide, TRBV19 and TRBV7-2 were expanded and the same genes were expanded by the CEF pool. The most frequent recombination observed was between TRBV19 and three TRBJ genes - TRBJ2-7, TRBJ2-3 and TRBJ1-5 (Figure 5F). Taken together, the results of CD8 T-cell activation and TCR expansion suggest that donor D384 was expanding GILG-specific CDR3s over non-GILG specific CDR3s, even when the GILG peptide was present along with five other HLA-A2 restricted immunodominant viral peptides in the CEF pool. This suggested that D384 immune repertoire was specifically sensitized to the influenza-A epitope.

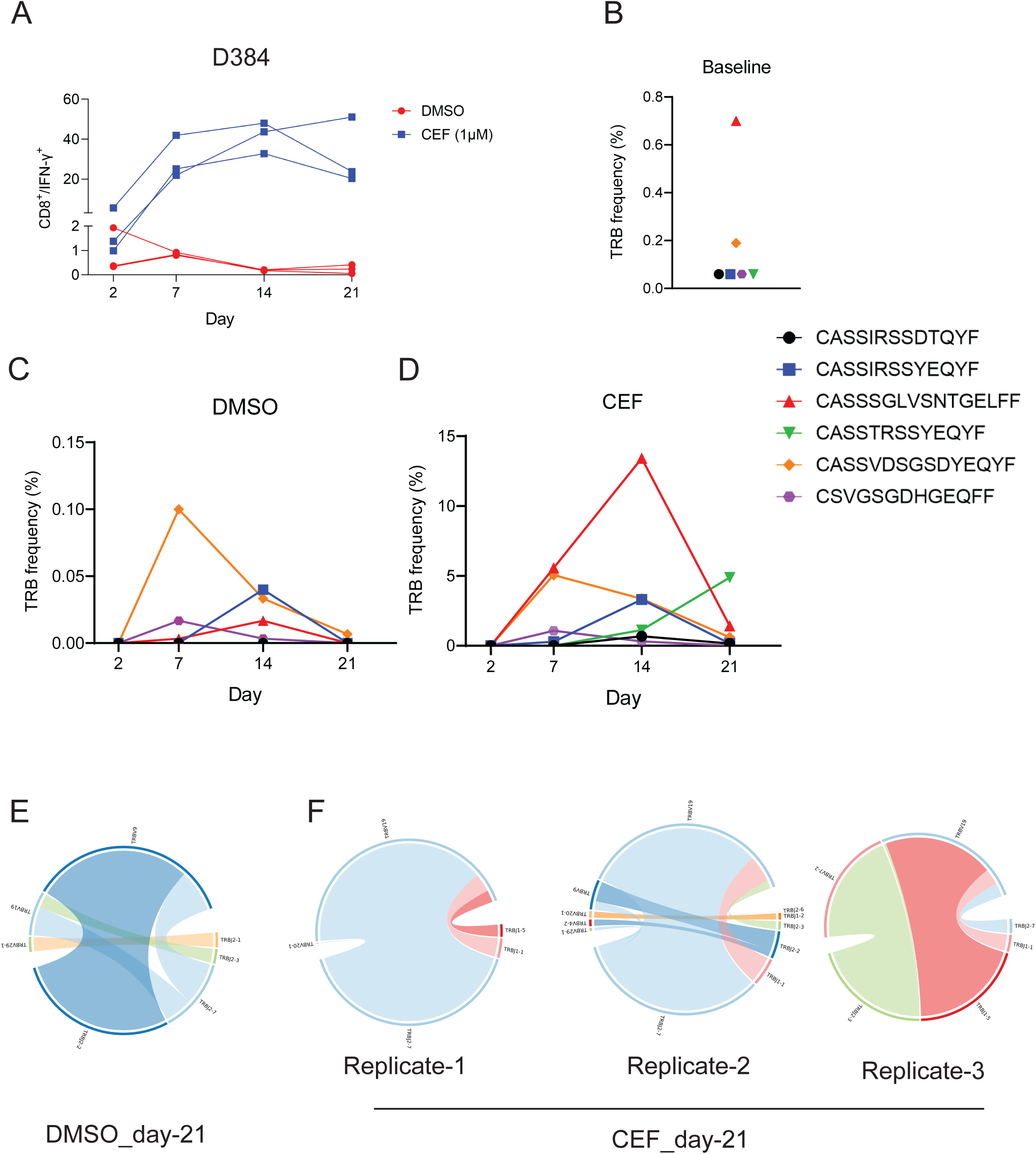

## Discussion

Influenza-A infection induces robust T-cell response conferring broad protection to a large fraction of the global population because of pre-existing viral immunity mediated by immunodominant epitopes encoded by the viral proteome. The HLA-A2 restricted GILGFVFTL peptide from influenza M protein has been well characterized as a 9-mer peptide in many donors. A large number of GILG-specific CDR3s shared between individuals (public TCR) is compiled in IEDB and VDJ databases.

We took advantage of this body of information to investigate the response of CD8 T-cells to the 9-mer peptide as well as a 15-mer version of the same epitope both in the context of T-cell activation as well as on the clonal expansion of antigen-specific T-cells captured by TCR sequencing.

The scope of this study was to obtain evidence that a 15-mer peptide carrying an immunodominant epitope is cross presented to CD8 T-cells and identified public CDR3s detected against the 9-mer GILGFVFTL peptide. Given that antigen-specific TCR discovery employs 15-mer overlapping peptides during screening, our study was intended to investigate if immunodominant epitopes will be missed by this approach. The findings from our study demonstrate that a 15-mer peptide carrying GILGFVFTL expanded many 9-mer epitope specific public TCRs. In addition, the 15-mer peptide also identified private CDR3s that were not expanded by the 9-mer peptide.

Garnering robust CD8 T-cell activation is dependent both on the peptide length and peptide dosage. Although, IFN-γ induction was rapid at 10μM 9-mer peptide, the response was not sustained compared to a 100-fold lower concentration of the nonameric peptide. In contrast, 15-mer peptide used at 10μM produced a robust and sustained response. The proportion of CD8 cells producing IFN-γ in the presence of 9-mer peptide was lower compared to the level of expansion of GILG-specific CDR3s suggesting that all expanded T-cells were not making IFN-γ. In addition, we observed expression of GZMB at higher antigen dose on day-21 suggesting apoptotic death associated with T-cell activation. as Activation induced cell death (AICD) is a a mechanism of maintaining tissue homeostasis by limiting the duration of T-cell activation, which is regulated by antigen dosage and affinity (27). Antigen dose and affinity shapes the T-cell repertoire by differentially regulating T-cell proliferation and differentiation into effector T-cells, the latter expressing IFN-γ (28, 29). Given that GILG peptide is a high affinity antigen our study shows that high dosage of the antigen can induce T-cells to become less responsive and lose their effector phenotype. In an in vitro setting, CTLs selected against the GILG epitope was found to have lower functional avidity (12), than what was observed in influenza-A infected subjects (30).

The top-10 GILG-specific CDR3s were expanded at different frequencies under conditions that induced robust CD8 T-cell activation and remained at a lower frequency under conditions that failed to activate T-cells. Similar to other studies, we detected, higher frequency of TRAV27/TRBV19 gene usage in activated T-cells whereas TRAV21, TRAV13.1/TRBV20.1, TRBV5.1 were enriched under conditions of minimal T-cell activation. TRAV27/TRBV19 TCRs are highly prevalent in donors and recognize GILG peptide and its naturally occurring variants (26, 31). The switch in TRAV and TRBV usage in activated T-cells demonstrate antigen specificity for the 15-mer peptide and confirms that it is correctly processed to present the 9-mer GILGFVFTL epitope.

Our study leads to the conclusion that 15-mer peptides are correctly processed in *in vitro* CD8 T-cell activation assay to generate the 9-mer immunodominant epitope. The expanded CDR3s include those that are also expanded by the 9-mer peptide, including additional CDR3s that may be derived from CD4 T-cells or T-cells recognizing novel epitopes generated from the 15-mer peptide. Therefore, use of 15-mer peptides in high throughput screening is unlikely to miss the existence of shorter antigen-specific immunodominant epitopes.

## Methods

### T cell activation assay

Donor PBMCs were obtained from commercial vendors for this study. PBMCs were thawed, counted and analyzed using the diagnostic panel of antibodies (Table S5). PBMCs were rested overnight in RPMI containing 10% human serum (Table S5). For T-cell activation assays, 750,000 PBMCs were incubated either with DMSO (negative control) or with 9-mer and 15-mer peptides in 0.5 ml RPMI (Gibco) +10% Human AB serum (Sigma) + 10 ng/ml IL-15 and 10 IU of IL-2 (Stemcell Technologies, Canada). The culture media was replenished every three days with fresh media containing 10 IU of IL-2 and 10 ng/ml IL-15. On days 2, 7, 14 and 21 of incubation, fresh peptides were added to the culture. For intracellular cytokine staining, cells were treated with Brefeldin A (BD Biosciences) for 5 hours, fixed and permeabilized using BD Lysis solution and Perm2 solutions respectively followed by staining with T-cell activation panel of antibodies (Table S5). Stained cells were analyzed in BD Accuri C6 Plus to detect the expression of activation marker IFN-γ and apoptosis marker GZMB on CD8 T cells. Data was analyzed using FlowJo and % of CD8^+^/IFN-γ cells was quantitated. GraphPad Prizm 9 was used to generate the plots.

### TCR sequencing and data analysis

200,000 PBMCS was removed after 2, 7, 14 and 21 days from the T-cell activation assay and processed for bulk TCR sequencing.

### Bulk TCR sequencing

TCR repertoire profiling was performed using the SMARTer TCR α/β Profiling Kit (Takara Bio, USA) according to the manufacturer’s protocol. RNA was isolated using the Qiagen RNA isolation kit. 10ng RNA from antigen-induced PBMCs were used as the starting material. The kit uses SMART technology (<u>S</u>witching <u>M</u>echanism <u>A</u>t 5’end of <u>R</u>NA <u>T</u>emplate) with 5’RACE to capture the entire V(D)J variable regions of TCR transcripts followed by two rounds of semi-nested PCR to obtain TCR-α and the β-chain. Libraries are prepared analyzed for quality and quantity. Sequencing is performed using the 2*300 MiSeq Reagent Kit v3 (Illumina, Inc.).

## References

1. Welsh RM, Fujinami RS. Pathogenic epitopes, heterologous immunity and vaccine design. Nat Rev Microbiol. 2007;5(7):555–63.

2. Sahin U, Tureci O. Personalized vaccines for cancer immunotherapy. Science. 2018;359(6382):1355–60.

3. Jou J, Harrington KJ, Zocca MB, Ehrnrooth E, Cohen EEW. The Changing Landscape of Therapeutic Cancer Vaccines-Novel Platforms and Neoantigen Identification. Clin Cancer Res. 2021;27(3):689–703.

4. Sahin U, Oehm P, Derhovanessian E, Jabulowsky RA, Vormehr M, Gold M, et al. An RNA vaccine drives immunity in checkpoint-inhibitor-treated melanoma. Nature. 2020;585(7823):107–12.

5. Golob JL, Lugogo N, Lauring AS, Lok AS. SARS-CoV-2 vaccines: a triumph of science and collaboration. JCI Insight. 2021;6(9).

6. Buchy P, Badur S. Who and when to vaccinate against influenza. Int J Infect Dis. 2020;93:375–87.

7. Kern F, Surel IP, Brock C, Freistedt B, Radtke H, Scheffold A, et al. T-cell epitope mapping by flow cytometry. Nat Med. 1998;4(8):975–8.

8. Ritmahan W, Kesmir C, Vroomans RMA. Revealing factors determining immunodominant responses against dominant epitopes. Immunogenetics. 2020;72(1-2):109–18.

9. Billeskov R, Beikzadeh B, Berzofsky JA. The effect of antigen dose on T cell-targeting vaccine outcome. Hum Vaccin Immunother. 2019;15(2):407–11.

10. Sahay B, Nguyen CQ, Yamamoto JK. Conserved HIV Epitopes for an Effective HIV Vaccine. J Clin Cell Immunol. 2017;8(4).

11. Adland E, Hill M, Lavandier N, Csala A, Edwards A, Chen F, et al. Differential Immunodominance Hierarchy of CD8(+) T-Cell Responses in HLA-B*27:05- and -B*27:02-Mediated Control of HIV-1 Infection. J Virol. 2018;92(4).

12. Keskin DB, Reinhold BB, Zhang GL, Ivanov AR, Karger BL, Reinherz EL. Physical detection of influenza A epitopes identifies a stealth subset on human lung epithelium evading natural CD8 immunity. Proc Natl Acad Sci U S A. 2015;112(7):2151–6.

13. Ferretti AP, Kula T, Wang Y, Nguyen DMV, Weinheimer A, Dunlap GS, et al. Unbiased Screens Show CD8(+) T Cells of COVID-19 Patients Recognize Shared Epitopes in SARS-CoV-2 that Largely Reside outside the Spike Protein. Immunity. 2020;53(5):1095–107 e3.

14. Sharma G, Holt RA. T-cell epitope discovery technologies. Hum Immunol. 2014;75(6):514–9.

15. Grifoni A, Weiskopf D, Ramirez SI, Mateus J, Dan JM, Moderbacher CR, et al. Targets of T Cell Responses to SARS-CoV-2 Coronavirus in Humans with COVID-19 Disease and Unexposed Individuals. Cell. 2020;181(7):1489–501 e15.

16. Mateus J, Grifoni A, Tarke A, Sidney J, Ramirez SI, Dan JM, et al. Selective and cross-reactive SARS-CoV-2 T cell epitopes in unexposed humans. Science. 2020.

17. Kiecker F, Streitz M, Ay B, Cherepnev G, Volk HD, Volkmer-Engert R, et al. Analysis of antigen-specific T-cell responses with synthetic peptides--what kind of peptide for which purpose? Hum Immunol. 2004;65(5):523–36.

18. Moguche AO, Musvosvi M, Penn-Nicholson A, Plumlee CR, Mearns H, Geldenhuys H, et al. Antigen Availability Shapes T Cell Differentiation and Function during Tuberculosis. Cell Host Microbe. 2017;21(6):695–706 e5.

19. Clemmensen HS, Dube JY, McIntosh F, Rosenkrands I, Jungersen G, Aagaard C, et al. In Vivo Antigen Expression Regulates CD4 T Cell Differentiation and Vaccine Efficacy against Mycobacterium tuberculosis Infection. mBio. 2021;12(2).

20. Darrah PA, Patel DT, De Luca PM, Lindsay RW, Davey DF, Flynn BJ, et al. Multifunctional TH1 cells define a correlate of vaccine-mediated protection against Leishmania major. Nat Med. 2007;13(7):843–50.

21. Billeskov R, Lindenstrom T, Woodworth J, Vilaplana C, Cardona PJ, Cassidy JP, et al. High Antigen Dose Is Detrimental to Post-Exposure Vaccine Protection against Tuberculosis. Front Immunol. 2017;8:1973.

22. Nandakumar S, Kannanganat S, Posey JE, Amara RR, Sable SB. Attrition of T-cell functions and simultaneous upregulation of inhibitory markers correspond with the waning of BCG-induced protection against tuberculosis in mice. PLoS One. 2014;9(11):e113951.

23. Poon MML, Byington E, Meng W, Kubota M, Matsumoto R, Grifoni A, et al. Heterogeneity of human anti-viral immunity shaped by virus, tissue, age, and sex. Cell Rep. 2021;37(9):110071.

24. Pappu BP, Shrikant PA. Alteration of cell surface sialylation regulates antigen-induced naive CD8+ T cell responses. J Immunol. 2004;173(1):275–84.

25. Sobek V, Balkow S, Korner H, Simon MM. Antigen-induced cell death of T effector cells in vitro proceeds via the Fas pathway, requires endogenous interferon-gamma and is independent of perforin and granzymes. Eur J Immunol. 2002;32(9):2490–9.

26. Sant S, Grzelak L, Wang Z, Pizzolla A, Koutsakos M, Crowe J, et al. Single-Cell Approach to Influenza-Specific CD8(+) T Cell Receptor Repertoires Across Different Age Groups, Tissues, and Following Influenza Virus Infection. Front Immunol. 2018;9:1453.

27. Green DR, Droin N, Pinkoski M. Activation-induced cell death in T cells. Immunol Rev. 2003;193:70–81.

28. Anthony DD, Milkovich KA, Zhang W, Rodriguez B, Yonkers NL, Tary-Lehmann M, et al. Dissecting the T Cell Response: Proliferation Assays vs. Cytokine Signatures by ELISPOT. Cells. 2012;1(2):127–40.

29. Balyan R, Gund R, Ebenezer C, Khalsa JK, Verghese DA, Krishnamurthy T, et al. Modulation of Naive CD8 T Cell Response Features by Ligand Density, Affinity, and Continued Signaling via Internalized TCRs. J Immunol. 2017;198(5):1823–37.

30. Boon AC, de Mutsert G, Fouchier RA, Osterhaus AD, Rimmelzwaan GF. The hypervariable immunodominant NP418-426 epitope from the influenza A virus nucleoprotein is recognized by cytotoxic T lymphocytes with high functional avidity. J Virol. 2006;80(12):6024–32.

31. Valkenburg SA, Josephs TM, Clemens EB, Grant EJ, Nguyen TH, Wang GC, et al. Molecular basis for universal HLA-A*0201-restricted CD8+ T-cell immunity against influenza viruses. Proc Natl Acad Sci U S A. 2016;113(16):4440–5.

