## Supplemental Figures for "Immunodominant influenza epitope GILGFVFTL engage common and divergent TCRs when presented as a 9-mer or a 15-mer peptide"

A

| | Day-7 | Day-14 | Day-21 |
|---|---|---|---|
| | CD8$^+$/IFN-γ$^+$ | | |
| DMSO | 1.7 | 1.2 | 1.2 |
| DMSO | 1.1 | 1 | 0.2 |
| DMSO | 1 | 0.5 | 0.1 |
| 9-mer 10μM | 8.8 | 5.1 | 0.4 |
| 9-mer 10μM | 9.8 | 4.6 | 0.8 |
| 9-mer 10μM | 8.6 | 2.9 | 1.2 |
| 9-mer 100nM | 15.1 | 30 | 1.9 |
| 9-mer 100nM | 18.9 | 35.7 | 2.4 |
| 9-mer 100nM | 15.8 | 37.8 | 4.3 |
| 15-mer 10μM | 2.4 | 35.6 | 7.9 |
| 15-mer 10μM | 1.7 | 35.8 | 12 |
| 15-mer 10μM | 2.2 | 36.2 | 7.7 |
| 15-mer 100nM | 2.1 | 0.4 | 0.7 |
| 15-mer 100nM | 0.8 | 0.4 | 0.3 |
| 15-mer 100nM | 0.9 | 0.4 | 0.2 |

B

| | 48h | 7-day | 14-day | 21-day |
|---|---|---|---|---|
| | CD8 % (absolute) | | | |
| DMSO | 11 | 44 | 17 | 26 |
| DMSO | 12 | 40 | 21 | 16 |
| DMSO | 12 | 42 | 22 | 28 |
| 9-mer 10μM | 11 | 45 | 21 | 4 |
| 9-mer 10μM | 12 | 47 | 21 | 4 |
| 9-mer 10μM | 12 | 46 | 21 | 4 |
| 9-mer 100nM | 13 | 29 | 53 | 13 |
| 9-mer 100nM | 13 | 32 | 59 | 15 |
| 9-mer 100nM | 12 | 26 | 56 | 21 |
| 15-mer 10μM | 12 | 33 | 24 | 11 |
| 15-mer 10μM | 13 | 24 | 26 | 12 |
| 15-mer 10μM | 13 | 31 | 28 | 10 |
| 15-mer 100nM | 11 | 34 | 18 | 28 |
| 15-mer 100nM | 11 | 24 | 10 | 13 |
| 15-mer 100nM | 12 | 22 | 10 | 15 |

C

| | Day-7 | Day-14 | Day-21 |
|---|---|---|---|
| | CD8$^+$/GZMB$^+$ | | |
| DMSO | 0.6 | 0.7 | 0.9 |
| DMSO | 0.5 | 1 | 0.8 |
| DMSO | 1.3 | 1.5 | 0.7 |
| 9-mer 10μM | 0.2 | 0.7 | 3.7 |
| 9-mer 10μM | 0.8 | 0.7 | 3.4 |
| 9-mer 10μM | 0.4 | 0.8 | 1.8 |
| 9-mer 100nM | 0.4 | 0.1 | 0.7 |
| 9-mer 100nM | 0.5 | 0.2 | 0.8 |
| 9-mer 100nM | 0.7 | 0.1 | 0.2 |
| 15-mer 10μM | 0.4 | 0.6 | 1 |
| 15-mer 10μM | 0.3 | 0.5 | 0.7 |
| 15-mer 10μM | 0.2 | 1 | 0.3 |
| 15-mer 100nM | 1.5 | 0.9 | 0.3 |
| 15-mer 100nM | 1.9 | 2 | 0.5 |
| 15-mer 100nM | 0.7 | 0.7 | 0.3 |

Figure S1

A

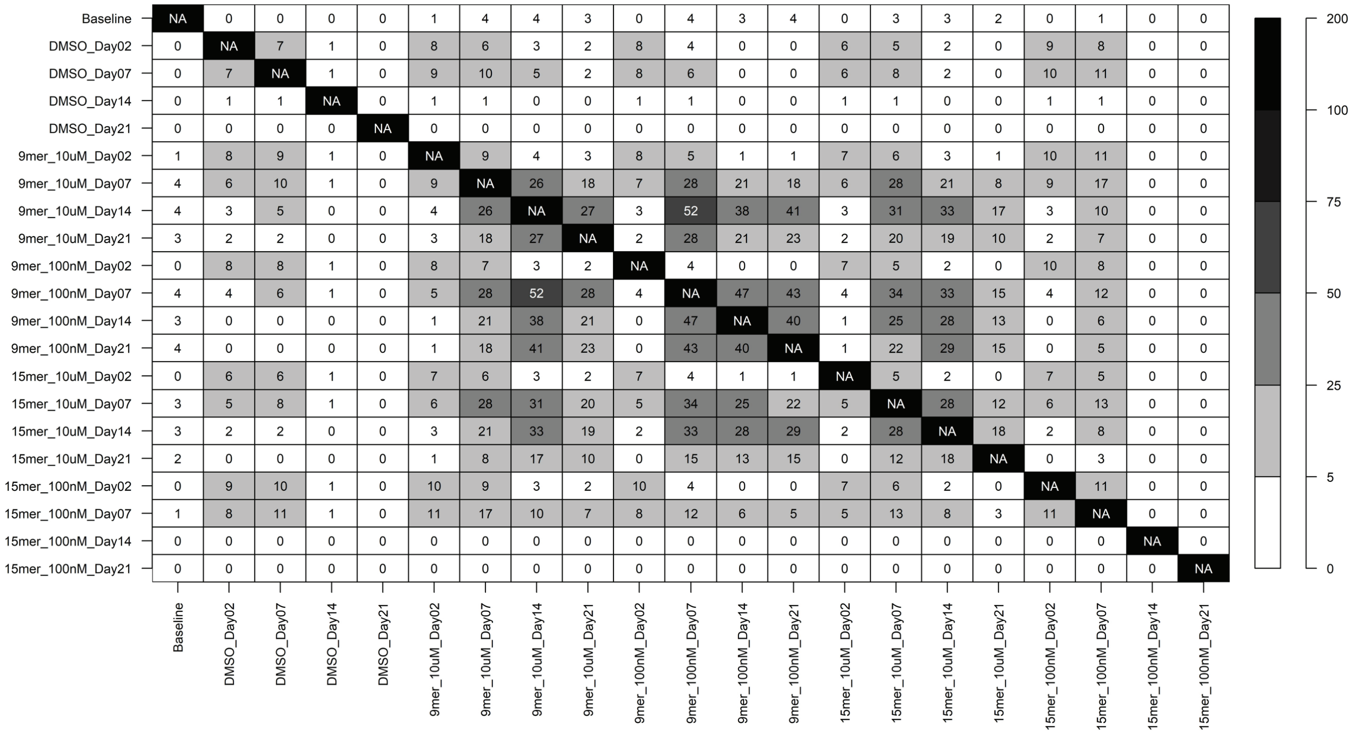

Overlap of GILG-specific CDR3

| | Baseline | DMSO_Day02 | DMSO_Day07 | DMSO_Day14 | DMSO_Day21 | 9mer_10uM_Day02 | 9mer_10uM_Day07 | 9mer_10uM_Day14 | 9mer_10uM_Day21 | 9mer_100nM_Day02 | 9mer_100nM_Day07 | 9mer_100nM_Day14 | 9mer_100nM_Day21 | 15mer_10uM_Day02 | 15mer_10uM_Day07 | 15mer_10uM_Day14 | 15mer_10uM_Day21 | 15mer_100nM_Day02 | 15mer_100nM_Day07 | 15mer_100nM_Day14 | 15mer_100nM_Day21 |
|---|---|---|---|---|---|---|---|---|---|---|---|---|---|---|---|---|---|---|---|---|---|
| Baseline | NA | 0 | 0 | 0 | 0 | 1 | 4 | 4 | 3 | 0 | 4 | 3 | 4 | 0 | 3 | 3 | 2 | 0 | 1 | 0 | 0 |
| DMSO_Day02 | 0 | NA | 7 | 1 | 0 | 8 | 6 | 3 | 2 | 8 | 4 | 0 | 0 | 6 | 5 | 2 | 0 | 9 | 8 | 0 | 0 |
| DMSO_Day07 | 0 | 7 | NA | 1 | 0 | 9 | 10 | 5 | 2 | 8 | 6 | 0 | 0 | 6 | 8 | 2 | 0 | 10 | 11 | 0 | 0 |
| DMSO_Day14 | 0 | 1 | 1 | NA | 0 | 1 | 1 | 0 | 0 | 1 | 1 | 0 | 0 | 1 | 1 | 0 | 0 | 1 | 1 | 0 | 0 |
| DMSO_Day21 | 0 | 0 | 0 | 0 | NA | 0 | 0 | 0 | 0 | 0 | 0 | 0 | 0 | 0 | 0 | 0 | 0 | 0 | 0 | 0 | 0 |
| 9mer_10uM_Day02 | 1 | 8 | 9 | 1 | 0 | NA | 9 | 4 | 3 | 8 | 5 | 1 | 1 | 7 | 6 | 3 | 1 | 10 | 11 | 0 | 0 |
| 9mer_10uM_Day07 | 4 | 6 | 10 | 1 | 0 | 9 | NA | 26 | 18 | 7 | 28 | 21 | 18 | 6 | 28 | 21 | 8 | 9 | 17 | 0 | 0 |
| 9mer_10uM_Day14 | 4 | 3 | 5 | 0 | 0 | 4 | 26 | NA | 27 | 3 | 52 | 38 | 41 | 3 | 31 | 33 | 17 | 3 | 10 | 0 | 0 |
| 9mer_10uM_Day21 | 3 | 2 | 2 | 0 | 0 | 3 | 18 | 27 | NA | 2 | 28 | 21 | 23 | 2 | 20 | 19 | 10 | 2 | 7 | 0 | 0 |
| 9mer_100nM_Day02 | 0 | 8 | 8 | 1 | 0 | 8 | 7 | 3 | 2 | NA | 4 | 0 | 0 | 7 | 5 | 2 | 0 | 10 | 8 | 0 | 0 |
| 9mer_100nM_Day07 | 4 | 4 | 6 | 1 | 0 | 5 | 28 | 52 | 28 | 4 | NA | 47 | 43 | 4 | 34 | 33 | 15 | 4 | 12 | 0 | 0 |
| 9mer_100nM_Day14 | 3 | 0 | 0 | 0 | 0 | 1 | 21 | 38 | 21 | 0 | 47 | NA | 40 | 1 | 25 | 28 | 13 | 0 | 6 | 0 | 0 |
| 9mer_100nM_Day21 | 4 | 0 | 0 | 0 | 0 | 1 | 18 | 41 | 23 | 0 | 43 | 40 | NA | 1 | 22 | 29 | 15 | 0 | 5 | 0 | 0 |
| 15mer_10uM_Day02 | 0 | 6 | 6 | 1 | 0 | 7 | 6 | 3 | 2 | 7 | 4 | 1 | 1 | NA | 5 | 2 | 0 | 7 | 5 | 0 | 0 |
| 15mer_10uM_Day07 | 3 | 5 | 8 | 1 | 0 | 6 | 28 | 31 | 20 | 5 | 34 | 25 | 22 | 5 | NA | 28 | 12 | 6 | 13 | 0 | 0 |
| 15mer_10uM_Day14 | 3 | 2 | 2 | 0 | 0 | 3 | 21 | 33 | 19 | 2 | 33 | 28 | 29 | 2 | 28 | NA | 18 | 2 | 8 | 0 | 0 |
| 15mer_10uM_Day21 | 2 | 0 | 0 | 0 | 0 | 1 | 8 | 17 | 10 | 0 | 15 | 13 | 15 | 0 | 12 | 18 | NA | 0 | 3 | 0 | 0 |
| 15mer_100nM_Day02 | 0 | 9 | 10 | 1 | 0 | 10 | 9 | 3 | 2 | 10 | 4 | 0 | 0 | 7 | 6 | 2 | 0 | NA | 11 | 0 | 0 |
| 15mer_100nM_Day07 | 1 | 8 | 11 | 1 | 0 | 11 | 17 | 10 | 7 | 8 | 12 | 6 | 5 | 5 | 13 | 8 | 3 | 11 | NA | 0 | 0 |
| 15mer_100nM_Day14 | 0 | 0 | 0 | 0 | 0 | 0 | 0 | 0 | 0 | 0 | 0 | 0 | 0 | 0 | 0 | 0 | 0 | 0 | 0 | NA | 0 |
| 15mer_100nM_Day21 | 0 | 0 | 0 | 0 | 0 | 0 | 0 | 0 | 0 | 0 | 0 | 0 | 0 | 0 | 0 | 0 | 0 | 0 | 0 | 0 | NA |

200
100
75
50
25
5
0

B

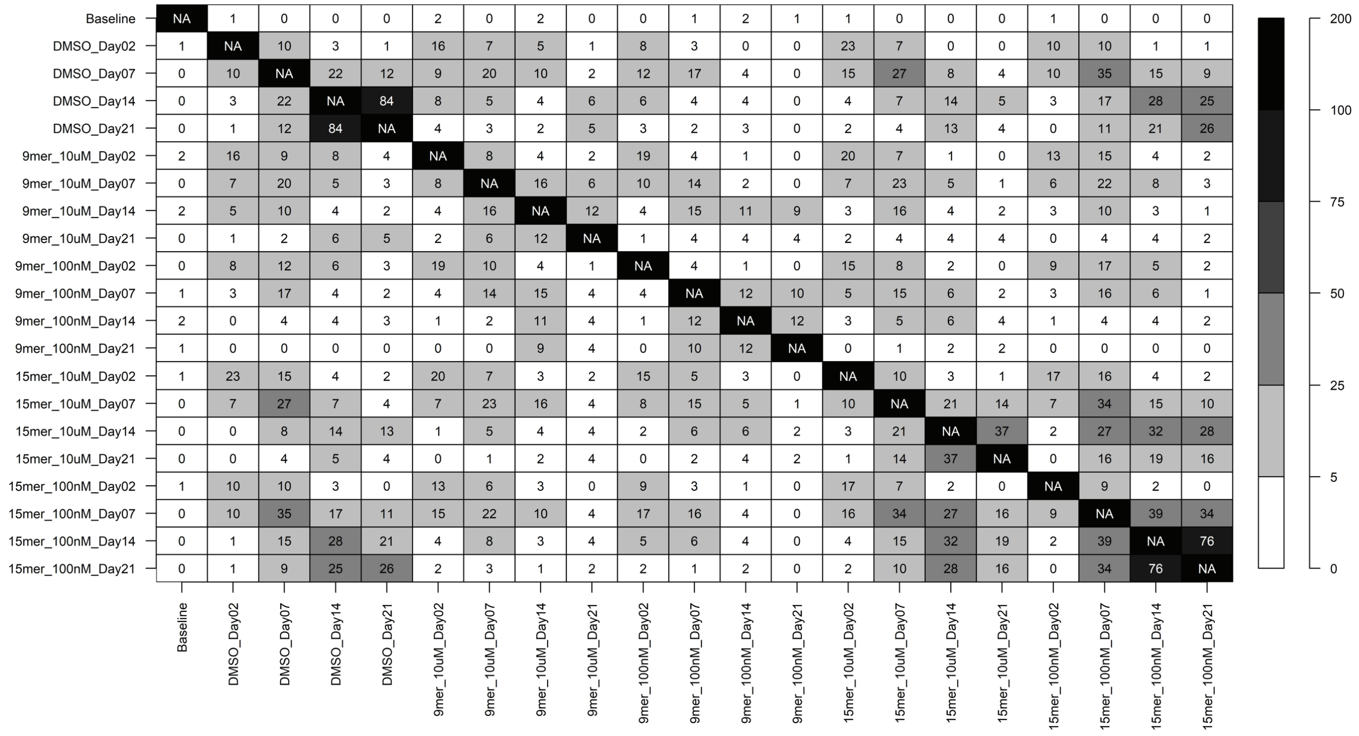

Non-GILG-specific CDR3

| | Baseline | DMSO_Day02 | DMSO_Day07 | DMSO_Day14 | DMSO_Day21 | 9mer_10uM_Day02 | 9mer_10uM_Day07 | 9mer_10uM_Day14 | 9mer_10uM_Day21 | 9mer_100nM_Day02 | 9mer_100nM_Day07 | 9mer_100nM_Day14 | 9mer_100nM_Day21 | 15mer_10uM_Day02 | 15mer_10uM_Day07 | 15mer_10uM_Day14 | 15mer_10uM_Day21 | 15mer_100nM_Day02 | 15mer_100nM_Day07 | 15mer_100nM_Day14 | 15mer_100nM_Day21 |
|---|---|---|---|---|---|---|---|---|---|---|---|---|---|---|---|---|---|---|---|---|---|
| Baseline | NA | 1 | 0 | 0 | 0 | 2 | 0 | 2 | 0 | 0 | 1 | 2 | 1 | 1 | 0 | 0 | 0 | 1 | 0 | 0 | 0 |
| DMSO_Day02 | 1 | NA | 10 | 3 | 1 | 16 | 7 | 5 | 1 | 8 | 3 | 0 | 0 | 23 | 7 | 0 | 0 | 10 | 10 | 1 | 1 |
| DMSO_Day07 | 0 | 10 | NA | 22 | 12 | 9 | 20 | 10 | 2 | 12 | 17 | 4 | 0 | 15 | 27 | 8 | 4 | 10 | 35 | 15 | 9 |
| DMSO_Day14 | 0 | 3 | 22 | NA | 84 | 8 | 5 | 4 | 6 | 6 | 4 | 4 | 0 | 4 | 7 | 14 | 5 | 3 | 17 | 28 | 25 |
| DMSO_Day21 | 0 | 1 | 12 | 84 | NA | 4 | 3 | 2 | 5 | 3 | 2 | 3 | 0 | 2 | 4 | 13 | 4 | 0 | 11 | 21 | 26 |
| 9mer_10uM_Day02 | 2 | 16 | 9 | 8 | 4 | NA | 8 | 4 | 2 | 19 | 4 | 1 | 0 | 20 | 7 | 1 | 0 | 13 | 15 | 4 | 2 |
| 9mer_10uM_Day07 | 0 | 7 | 20 | 5 | 3 | 8 | NA | 16 | 6 | 10 | 14 | 2 | 0 | 7 | 23 | 5 | 1 | 6 | 22 | 8 | 3 |
| 9mer_10uM_Day14 | 2 | 5 | 10 | 4 | 2 | 4 | 16 | NA | 12 | 4 | 15 | 11 | 9 | 3 | 16 | 4 | 2 | 3 | 10 | 3 | 1 |
| 9mer_10uM_Day21 | 0 | 1 | 2 | 6 | 5 | 2 | 6 | 12 | NA | 1 | 4 | 4 | 4 | 2 | 4 | 4 | 4 | 0 | 4 | 4 | 2 |
| 9mer_100nM_Day02 | 0 | 8 | 12 | 6 | 3 | 19 | 10 | 4 | 1 | NA | 4 | 1 | 0 | 15 | 8 | 2 | 0 | 9 | 17 | 5 | 2 |
| 9mer_100nM_Day07 | 1 | 3 | 17 | 4 | 2 | 4 | 14 | 15 | 4 | 4 | NA | 12 | 10 | 5 | 15 | 6 | 2 | 3 | 16 | 6 | 1 |
| 9mer_100nM_Day14 | 2 | 0 | 4 | 4 | 3 | 1 | 2 | 11 | 4 | 1 | 12 | NA | 12 | 3 | 5 | 6 | 4 | 1 | 4 | 4 | 2 |
| 9mer_100nM_Day21 | 1 | 0 | 0 | 0 | 0 | 0 | 0 | 9 | 4 | 0 | 10 | 12 | NA | 0 | 1 | 2 | 2 | 0 | 0 | 0 | 0 |
| 15mer_10uM_Day02 | 1 | 23 | 15 | 4 | 2 | 20 | 7 | 3 | 2 | 15 | 5 | 3 | 0 | NA | 10 | 3 | 1 | 17 | 16 | 4 | 2 |
| 15mer_10uM_Day07 | 0 | 7 | 27 | 7 | 4 | 7 | 23 | 16 | 4 | 8 | 15 | 5 | 1 | 10 | NA | 21 | 14 | 7 | 34 | 15 | 10 |
| 15mer_10uM_Day14 | 0 | 0 | 8 | 14 | 13 | 1 | 5 | 4 | 4 | 2 | 6 | 6 | 2 | 3 | 21 | NA | 37 | 2 | 27 | 32 | 28 |
| 15mer_10uM_Day21 | 0 | 0 | 4 | 5 | 4 | 0 | 1 | 2 | 4 | 0 | 2 | 4 | 2 | 1 | 14 | 37 | NA | 0 | 16 | 19 | 16 |
| 15mer_100nM_Day02 | 1 | 10 | 10 | 3 | 0 | 13 | 6 | 3 | 0 | 9 | 3 | 1 | 0 | 17 | 7 | 2 | 0 | NA | 9 | 2 | 0 |
| 15mer_100nM_Day07 | 0 | 10 | 35 | 17 | 11 | 15 | 22 | 10 | 4 | 17 | 16 | 4 | 0 | 16 | 34 | 27 | 16 | 9 | NA | 39 | 34 |
| 15mer_100nM_Day14 | 0 | 1 | 15 | 28 | 21 | 4 | 8 | 3 | 4 | 5 | 6 | 4 | 0 | 4 | 15 | 32 | 19 | 2 | 39 | NA | 76 |
| 15mer_100nM_Day21 | 0 | 1 | 9 | 25 | 26 | 2 | 3 | 1 | 2 | 2 | 1 | 2 | 0 | 2 | 10 | 28 | 16 | 0 | 34 | 76 | NA |

200
100
75
50
25
5
0

Figure S2

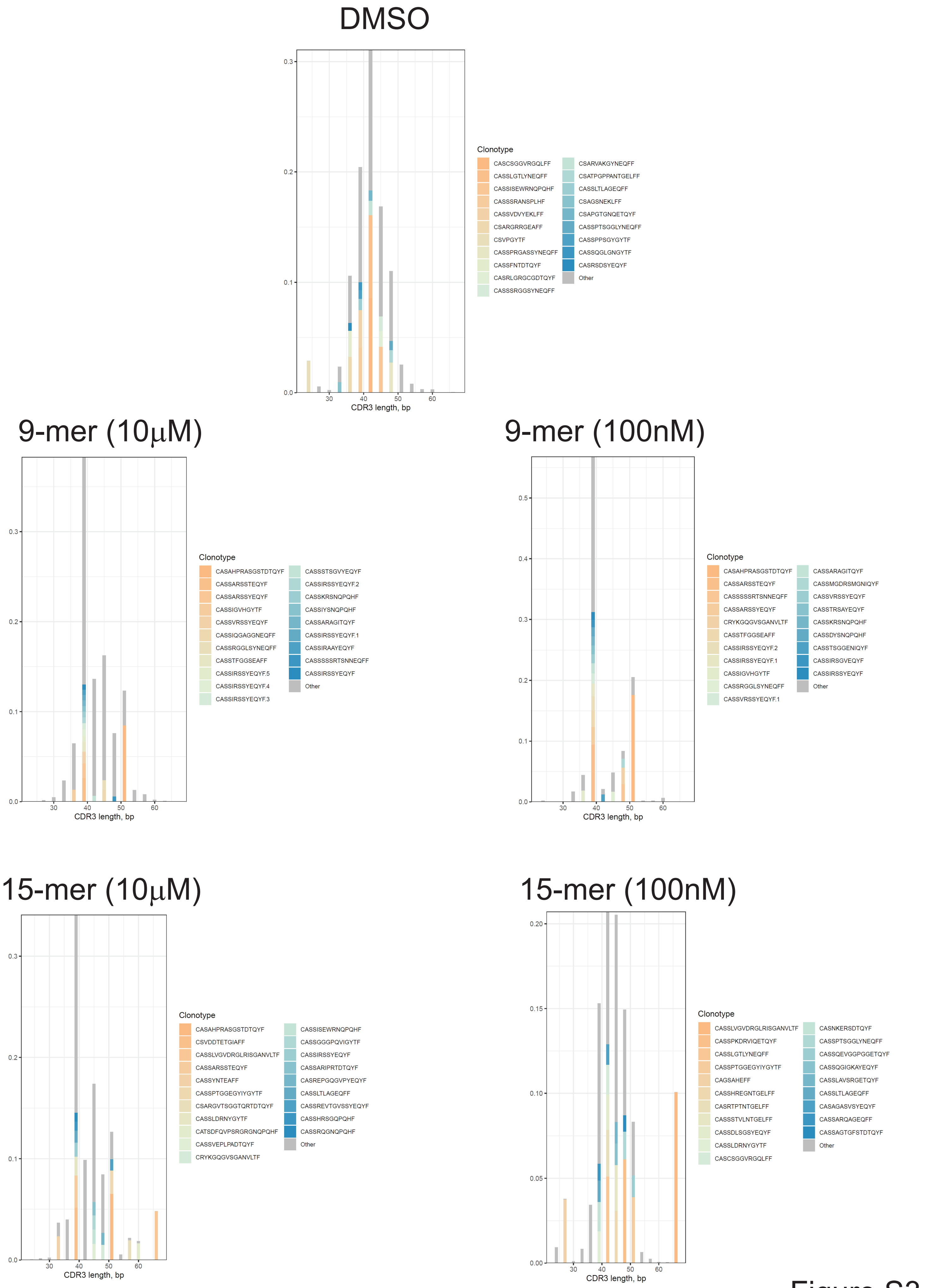
DMSO
CDR3 length, bp
Clonotype
CASCSGGVRGQLFF
CASSLGTLYNEQFF
CASSISEWRNQPQHF
CASSSRANSPLHF
CASSVDVYEKLFF
CSARGRRGEAFF
CSVPGYTF
CASSPRGASSYNEQFF
CASSFNTDTQYF
CASRLGRGCGDTQYF
CASSSRGGSYNEQFF
CSARVAKGYNEQFF
CSATPGPPANTGELFF
CASSLTLAGEQFF
CSAGSNEKLFF
CSAPGTGNQETQYF
CASSPTSGGLYNEQFF
CASSPPSGYGYTF
CASSQGLGNGYTF
CASRSDSYEQYF
Other
9-mer (10μM)
CDR3 length, bp
Clonotype
CASAHPRASGSTDTQYF
CASSARSSTEQYF
CASSARSSYEQYF
CASSIGVHGYTF
CASSVRSSYEQYF
CASSIQGAGGNEQFF
CASSRGGLSYNEQFF
CASSTFGGSEAFF
CASSIRSSYEQYF.5
CASSIRSSYEQYF.4
CASSIRSSYEQYF.3
CASSSTSGVYEQYF
CASSIRSSYEQYF.2
CASSKRSNQPQHF
CASSIYSNQPQHF
CASSARAGITQYF
CASSIRSSYEQYF.1
CASSIRAAYEQYF
CASSSSSRTSNNEQFF
CASSIRSSYEQYF
Other
9-mer (100nM)
CDR3 length, bp
Clonotype
CASAHPRASGSTDTQYF
CASSARSSTEQYF
CASSSSSRTSNNEQFF
CASSARSSYEQYF
CRYKGQGVSGANVLTF
CASSTFGGSEAFF
CASSIRSSYEQYF.2
CASSIRSSYEQYF.1
CASSIGVHGYTF
CASSRGGLSYNEQFF
CASSVRSSYEQYF.1
CASSARAGITQYF
CASSMGDRSMGNIQYF
CASSVRSSYEQYF
CASSTRSAYEQYF
CASSKRSNQPQHF
CASSDYSNQPQHF
CASSTSGGENIQYF
CASSIRSGVEQYF
CASSIRSSYEQYF
Other
15-mer (10μM)
CDR3 length, bp
Clonotype
CASAHPRASGSTDTQYF
CSVDDTETGIAFF
CASSLVGVDRGLRISGANVLTF
CASSARSSTEQYF
CASSYNTEAFF
CASSPTGGEGYIYGYTF
CSARGVTSGGTQRTDTQYF
CASSLDRNYGYTF
CATSDFQVPSRGRGNQPQHF
CASSVEPLPADTQYF
CRYKGQGVSGANVLTF
CASSISEWRNQPQHF
CASSGGGPQVIGYTF
CASSIRSSYEQYF
CASSARIPRTDTQYF
CASREPGQGVPYEQYF
CASSLTLAGEQFF
CASSREVTGVSSYEQYF
CASSHRSGQPQHF
CASSRQGNQPQHF
Other
15-mer (100nM)
CDR3 length, bp
Clonotype
CASSLVGVDRGLRISGANVLTF
CASSPKDRVIQETQYF
CASSLGTLYNEQFF
CASSPTGGEGYIYGYTF
CAGSAHEFF
CASSHREGNTGELFF
CASRTPTNTGELFF
CASSSTVLNTGELFF
CASSDLSGSYEQYF
CASSLDRNYGYTF
CASCSGGVRGQLFF
CASNKERSDTQYF
CASSPTSGGLYNEQFF
CASSQEVGGPGGETQYF
CASSQGIGKAYEQYF
CASSLAVSRGETQYF
CASSLTLAGEQFF
CASAGASVSYEQYF
CASSARQAGEQFF
CASSAGTGFSTDTQYF
Other

Figure S3

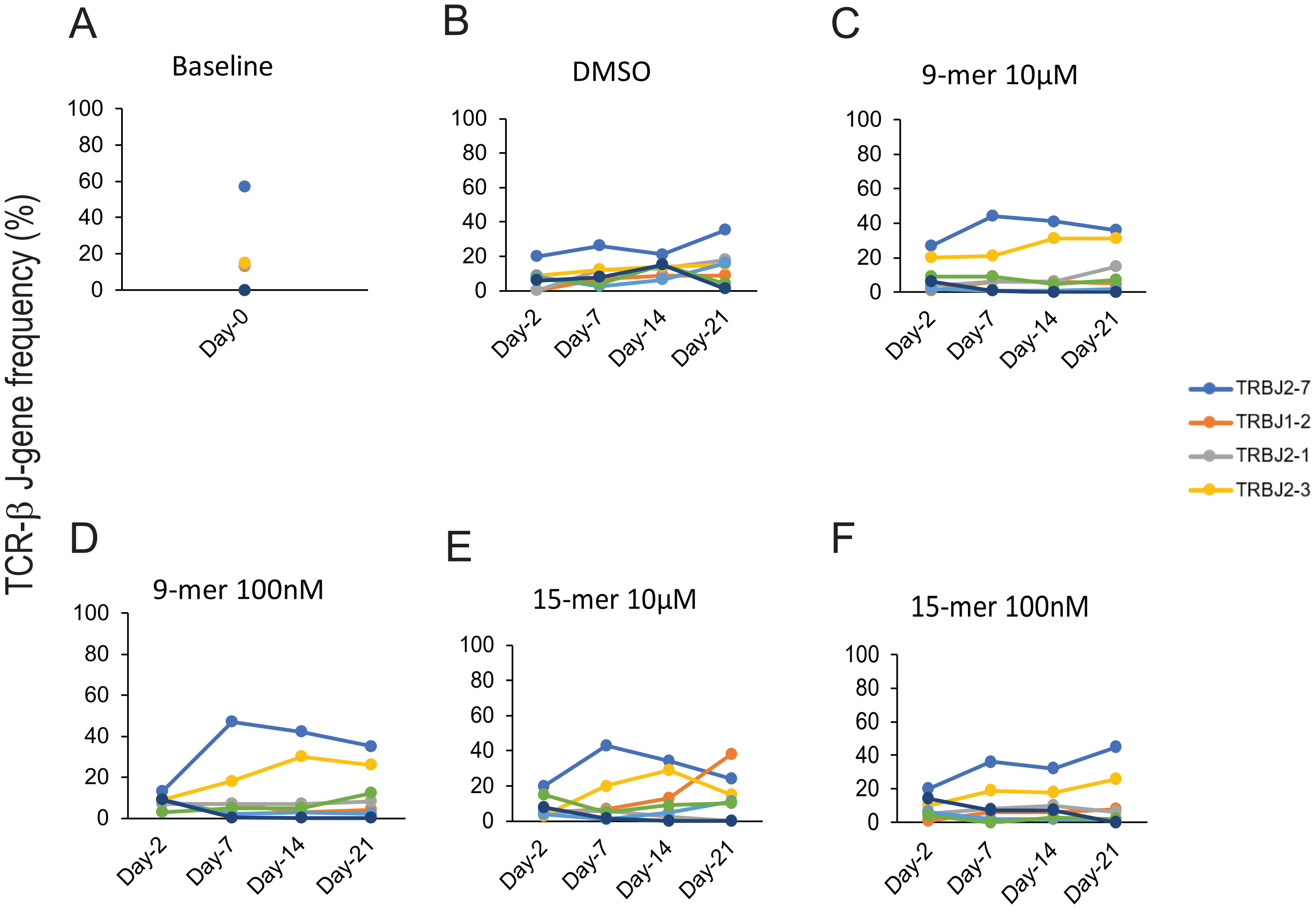

A
Baseline
100
80
60
40
20
0
Day-0
B
DMSO
100
80
60
40
20
0
Day-2
Day-7
Day-14
Day-21
C
9-mer 10µM
100
80
60
40
20
0
Day-2
Day-7
Day-14
Day-21
TRBJ2-7
TRBJ1-2
TRBJ2-1
TRBJ2-3
TCR-β J-gene frequency (%)
D
9-mer 100nM
100
80
60
40
20
0
Day-2
Day-7
Day-14
Day-21
E
15-mer 10µM
100
80
60
40
20
0
Day-2
Day-7
Day-14
Day-21
F
15-mer 100nM
100
80
60
40
20
0
Day-2
Day-7
Day-14
Day-21

Figure S4

### Non-GILG CDR3

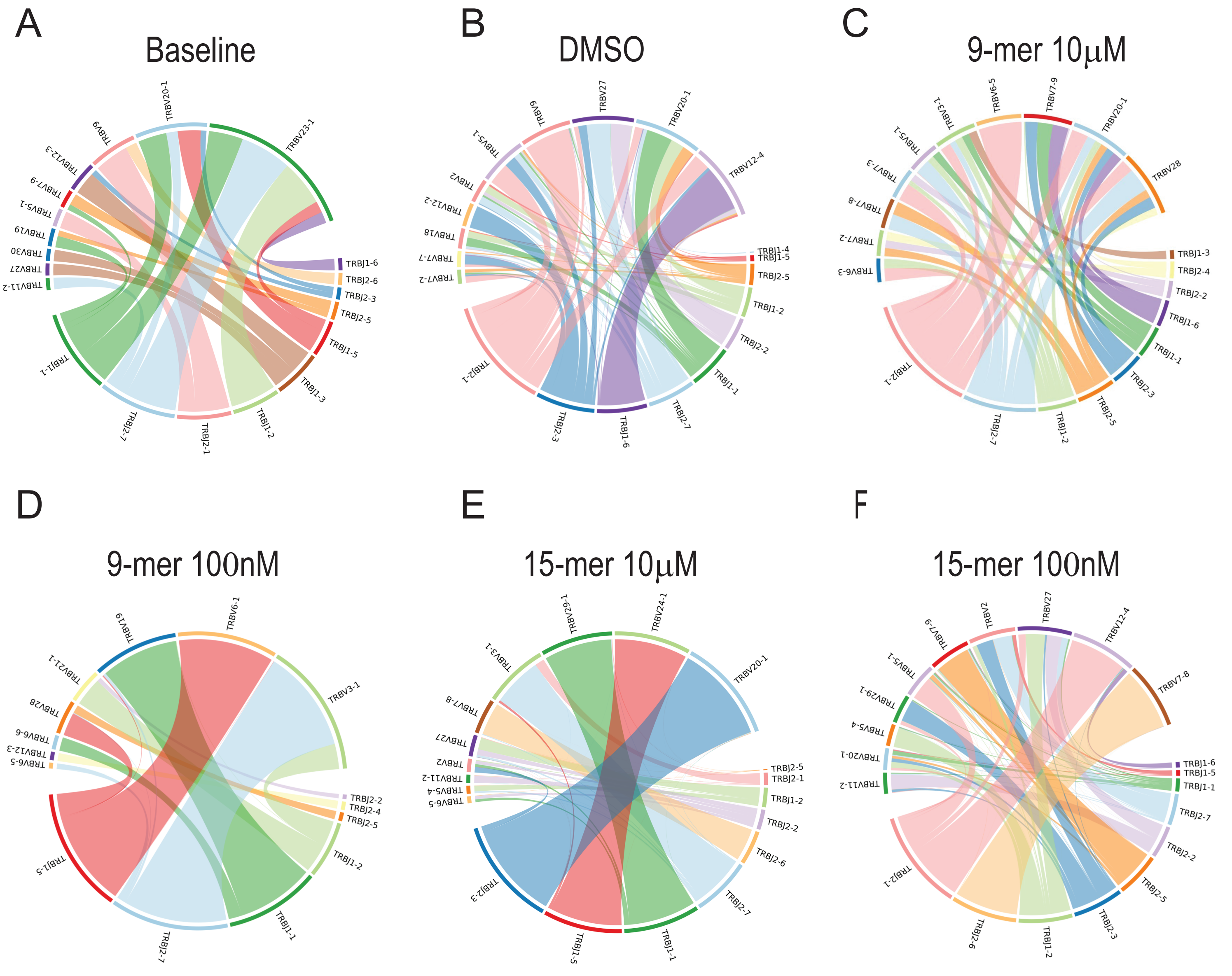

Figure S5
